# HPF1 dynamically controls the PARP1/2 balance between initiating and elongating ADP- ribose modifications

**DOI:** 10.1101/2021.05.19.444852

**Authors:** Marie-France Langelier, Ramya Billur, Aleksandr Sverzhinsky, Ben E. Black, John M. Pascal

## Abstract

Upon detecting DNA strand breaks, PARP1 and PARP2 produce the posttranslational modification poly(ADP-ribose) to orchestrate the cellular response to DNA damage. Histone PARylation factor 1 (HPF1) binds to PARP1/2 to directly regulate their catalytic output. HPF1 is required for the modification of serine residues with ADP-ribose, whereas glutamate/aspartate residues are modified in the absence of HPF1. PARP1 is an abundant nuclear protein, whereas HPF1 is present in much lower amounts, raising the question of whether HPF1 can pervasively modulate PARP1 activity. Here we show biochemically that HPF1 efficiently regulates PARP1/2 catalytic output at the sub-stoichiometric ratios matching their relative cellular abundances. HPF1 rapidly associates and dissociates from multiple PARP1 molecules, initiating ADP-ribose modification of serine residues before modification can initiate on glutamate/aspartate residues. HPF1 accelerates the rate of attaching the first ADP-ribose, such that this initiation event is comparable to the rate of the elongation reaction to form poly(ADP-ribose). This “hit and run” mechanism ensures that HPF1 contributions to the PARP1/2 active site during initiation do not persist and interfere with PAR chain elongation at sites of DNA damage. HPF1 thereby balances initiation and elongation events to regulate PARP1/2 output. Structural analysis of HPF1 in complex with PARP1 provides first insights into the assembly on a DNA strand break, and the HPF1 impact on PARP1 retention on DNA. Our data support the prevalence of the serine-ADP-ribose modification in cells and establish that HPF1 imparts the efficiency of serine-ADP-ribose modification required for an acute response to DNA damage.

## INTRODUCTION

Genome integrity in eukaryotic cells is maintained by a variety of surveillance pathways. Poly(ADP-ribose) polymerase (PARP) enzymes 1, 2, and 3 are considered first responders in the cellular response to DNA damage since they rapidly detect and signal DNA strand breaks ^1^. PARP1/2/3 binding to DNA breaks strongly stimulates their catalytic activities, which consists of covalently attaching ADP-ribose to target proteins using NAD^+^ as a source of ADP-ribose. While PARP3 only adds mono ADP-ribose (MAR) to its targets, PARP1 and PARP2 can form chains of ADP-ribose units known as poly(ADP-ribose) or PAR ^2^. The catalytic domain structure of PARP1 (Fig. 1A) indicated an “acceptor” site where ADP-ribose can bind and be elongated with another ADP-ribose unit from the NAD^+^ “donor” site ^3^. PARP1/2/3 modify themselves with ADP-ribose at multiple sites, a process termed auto-modification, and they also modify a variety of other proteins including histones, other DNA repair factors, and certain DNA structures ^4–6^. The rapid production of PAR and MAR at and around the site of DNA damage allows recruitment of DNA repair factors harboring PAR and MAR binding domains to start the repair process. The ADP-ribose modification can also modulate the local structure of chromatin to facilitate repair, and is also proposed to seed the assembly of phase condensates that create a repair environment ^7^.

**Figure 1.**
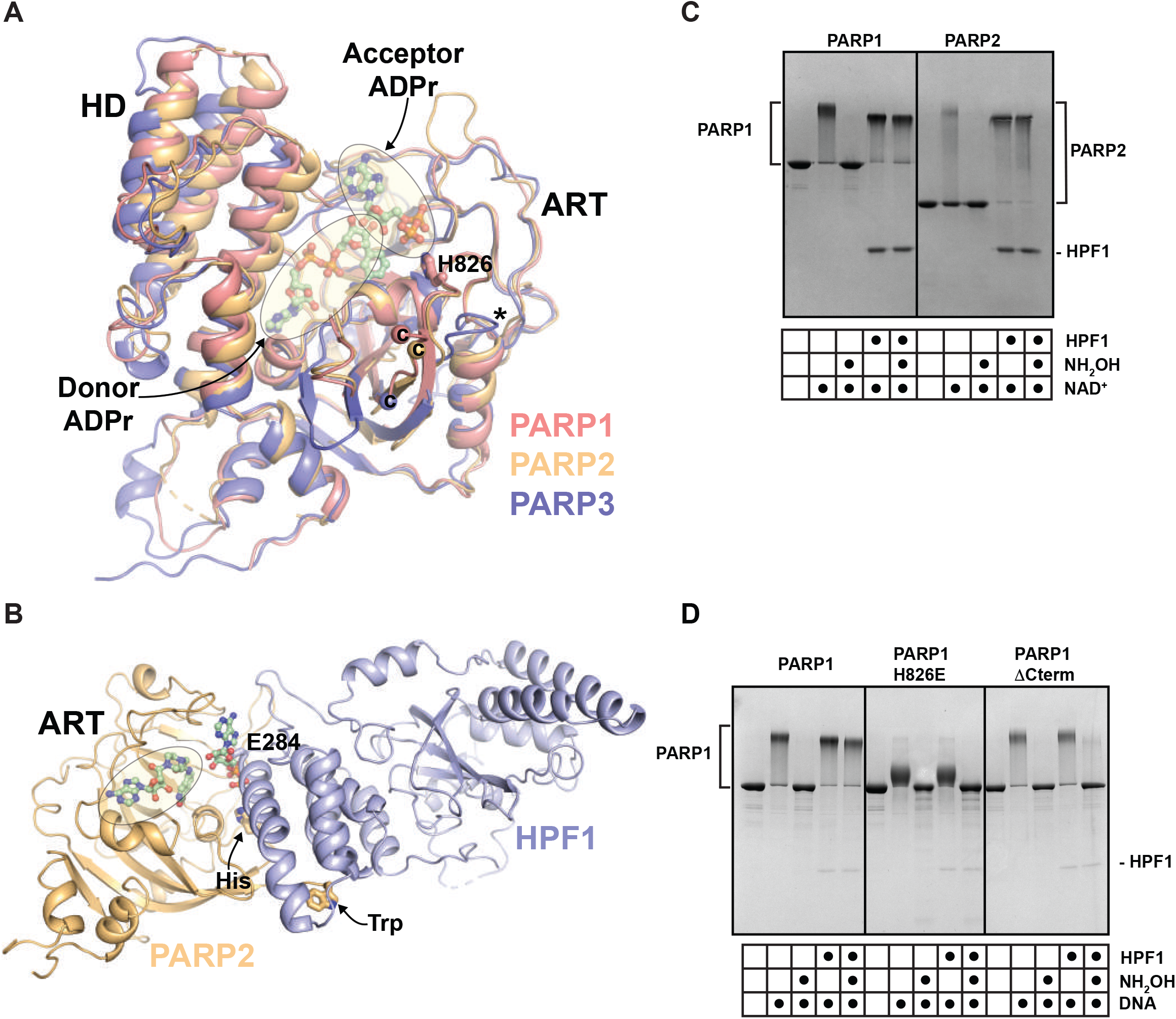
HPF1 regulation of PARP1 and PARP2. A) The catalytic domains of PARP1 (4dqy), PARP2 (4pjv), and PARP3 (3fhb) are superimposed. The helical domain (HD) and the ADP-ribosyltransferase (ART) domain are labeled. The Donor ADPr site is defined by the PARP1 complex with BAD (6bhv), and the Acceptor ADPr site is defined by the PARP1 complex with carba-NAD^+^ (1a26). PARP1 and PARP2 share a similar structure of the loop that bears residue H826 (PARP1), whereas PARP3 has a truncated loop (*). PARP1 and PARP2 share conserved residues at their C-terminal ends that are not conserved in PARP3 (C-termini labeled “C”). B) The ART of PARP2 (HD deleted) in complex with HPF1 (6tx3). A conserved His residue from the loop highlighted in panel B (826 in PARP1), and a Trp residue at the C-terminus (1014 in PARP1), are drawn as sticks and labeled (His and Trp). The Donor site is bound by the NAD^+^ mimic EB-47 that was crystallized with the complex. HPF1 binding clashes with the Acceptor site (carba-NAD^+^ is shown bound to the acceptor site as in panel B in order to highlight the clash). HPF1 residue E284 directly contributes to the PARP active site. C) PARP1 or PARP2 (1 μM) was incubated with dumbbell DNA containing a central nick (1 μM) with or without HPF1 (1 μM) for 10 min at room temperature (RT). 500 μM NAD^+^ was added for 5 min and reactions were quenched with 500 µM PARP inhibitor (olaparib or talazoparib). Where indicated, reactions were treated with 1M hydroxylamine (NH_2_OH) for one hour. Reactions were resolved by SDS-PAGE and treated with Imperial Stain. D) Same as panel C for PARP1 WT, mutant H826E, and mutant ΔCterm (Δ1012-1014), with 0.1 μM of HPF1.

PARP1 is a modular protein composed of six domains. Zinc binding domains Zn1 and Zn2 recognize and bind to the DNA break (also referred to as zinc finger domains FI and FII; ^8,9^. A third zinc-binding domain (Zn3, or FIII) and the Trp-Gly-Arg (WGR) domain contribute to DNA binding and support interdomain contacts that transmit the DNA binding signal to the catalytic domain (CAT) ^10^. The CAT domain is composed of two subdomains: the helical domain (HD) and the ADP-ribosyl transferase domain (ART). The HD regulates PARP1 activity by blocking NAD^+^ binding to the active site ^11^. A BRCT domain is located adjacent to an extended linker region that bears the primary sites for PARP1 automodification. When PARP1 binds to DNA damage, the assembly of domains and their interactions on DNA promote partial unfolding of the HD allowing access of NAD^+^ to the catalytic site ^10–12^. PARP2 and PARP3 domain structures are simpler, consisting of only a WGR and a CAT domain with short unstructured regions at their N-termini ^13–15^. The HD substrate-blocking mechanism, which is relieved by DNA binding, is conserved in PARP2 and PARP3 ^11,12^.

Histone PARylation Factor 1 (HPF1) is a central regulator of PARP1 and PARP2 activity in the DNA damage response ^16^. Indeed, HPF1 switches the amino acid specificity of PARP1 and PARP2 modifications from Glu/Asp to Ser ADP-ribosylation ^16^. HPF1 was also shown to reduce PARP1 catalytic output, to decrease the length of PAR formed by PARP1 and PARP2, and to promote *trans* ADP-ribosylation of histones in relation to *cis* modification of PARP1 itself ^17^. Crystal structures of HPF1 bound to the PARP2 CAT domain lacking an HD ^18^, or to the PARP1 CAT domain lacking an HD ^19^, have revealed the mechanistic basis for HPF1 effect on PARP1 and PARP2. HPF1 binds to PARP1 and PARP2 CAT and inserts a Glu residue to complement the active site (Fig. 1B). The HD limits HPF1 binding; therefore, unfolding of the HD through PARP1/2 interaction with DNA (or deleting the HD), is a pre-requisite for the most stable interaction with HPF1. The HPF1 interaction with PARP1/2 is further stabilized by compounds that maintain the HD in an open conformation, such as the NAD^+^ mimic known as EB-47 ^18^. By comparison to the active site of cholera toxin-like ADP-ribosyl transferases, PARP1 and PARP2 are deemed to lack a second Glu residue, in addition to the catalytic Glu of the conserved His-Tyr-Glu triad motif (Glu988 in PARP1 and Glu545 in PARP2) ^18^. The second Glu is provided by HPF1 and deprotonates Ser residues to allow their ADP-ribosylation ^18,19^. In contrast to Ser, Glu and Asp are naturally deprotonated at neutral pH and therefore do not require HPF1 to be ADP-ribosylated by PARP1. In further support, a recent cryo-electron microscopy (EM) structure of a PARP2-HPF1 complex bound to nucleosomes provided a snapshot of how PARP2 assembles with HPF1 on a DNA break in the context of chromatin ^20^. In each of the HPF1 complexes with PARP1/2, HPF1 sterically blocks the elongation or “acceptor” sites, thus indicating why HPF1 has a strong influence on PARP1/2 catalytic output and the distribution of PAR lengths produced.

Structural analysis has highlighted a 1:1 complex of HPF1 with PARP1/2 ready to perform ADP-ribose modification of Ser ^18,19^. Moreover, PARP1 modification of Ser residues is the predominant form of PARylation performed by PARP1 in response to DNA damage ^21^. However, HPF1 is about 20-fold less abundant than PARP1 in the cell ^16,22^, raising the question of how HPF1 could function efficiently to support Ser modification by PARP1. Our study reveals that HPF1, in fact, can work efficiently to support Ser modification over Glu/Asp modification even at sub-stoichiometric ratios relative to PARP1, and the regulation is aided by a dynamic interaction between HPF1 and PARP1 as indicated by surface plasmon resonance (SPR) analysis of the interaction. We find that HPF1 regulates PARP1 and PARP2 activity not only by blocking the “acceptor” site and restricting PAR elongation, but also by markedly stimulating the rate of initiation of ADP-ribosylation, such that it is more comparable to the rate of elongation. HPF1 thus regulates the balance between initiation and elongation of the ADP-ribose modification. We also present <u>h</u>ydrogen/deuterium e<u>x</u>change <u>m</u>ass <u>s</u>pectrometry (HXMS) data that support the dynamic PARP1/HPF1 interaction and indicate that HPF1 can further push the HD to the unfolded conformation induced by DNA damage binding, thereby contributing to PARP1 persistence on DNA damage. Negative stain EM analysis of the PARP1-HPF1 complex bound to a DNA single-strand break provide first insights into the overall assembly of this dynamic complex. Our results support a model where HPF1 permits PARP1/2 initiation on Ser residues at a rate that greatly exceeds the rate of initiation on Glu/Asp residues, dynamically engages multiple PARP1/2 molecules before Glu/Asp initiation events occur, and suppresses elongation and thereby increases the opportunities for further initiation events. Together, our results and model explain how HPF1 efficiently supports modification of Ser residues by PARP1 in the cell despite its low abundance compared to PARP1, and they provide new understanding of the dynamic regulation of ADP-ribosylation in response to DNA damage.

## RESULTS

HPF1 adapts the catalytic output of PARP1 and PARP2 such that ADP-ribose is linked to Ser residues rather than Glu/Asp residues ^16^. The switching effect can be easily visualized by SDS-PAGE analysis where covalent PAR attachment causes a pronounced shift in PARP1/2 protein migration (Fig. 1C). Whereas the ester bond of PAR linked to Glu/Asp residues is sensitive to hydroxylamine treatment and thus reverses the PARP1/2 migration shift, the ether bond of PAR linked to Ser residues, due to the presence of HPF1, is resistant to hydroxylamine treatment (Fig. 1C) ^16^. In contrast, PARP3 is not regulated by HPF1 and does not exhibit changes in hydroxylamine sensitivity (Supp. Fig. 1) ^16^. Prior to the structures of HPF1 bound to PARP1/PARP2 ^18–20^, we looked for structural features that might underlie the specificity toward PARP1/2, and we noted that PARP1/2 shared two features relative to PARP3 that localized to a similar region of the protein surface: a conserved Trp residue at the extreme C-terminus (PARP1 – Trp1014; PARP2 – Trp586), and a nearby loop containing a conserved His residue (PARP1 – His826; PARP2 – His381) (Fig. 1A). PARP1 bearing a three amino acid C-terminal deletion that removes Trp1014 (PARP1ΔCterm) was less responsive to HPF1, but otherwise showed wild-type levels of PAR production (Fig. 1D). Likewise, a charge reversal mutant of PARP1 at position His826 (H826E) was resistant to the effects of HPF1 on PAR linkage. His826 contributes to a region of PARP1 that regulates the PAR elongation reaction ^3^ (Fig. 1A), and the mutant H826E indeed exhibits deficiencies in PAR production, making polymers of short sizes as evidenced by a reduced migration shift on SDS-PAGE (Fig. 1D). Regardless, the PAR produced by PARP1 H826E is Glu/Asp-linked in both the presence and absence of HPF1. Recent NMR, crystallographic, and mutagenic analyses of PARP1/2 interaction with HPF1 demonstrated that HPF1 indeed engages these distinct features of PARP1 and PARP2 ^18,19^. Our results are thus consistent with these structural snapshots, and the PARP1 mutants that we generated serve as controls for modulating the HPF1 interaction.

In our biochemical analysis of HPF1 control of PARP1 catalytic output, we noted that HPF1 could efficiently regulate the PARP1 specificity switch to Ser residues at sub-stoichiometric ratios, maintaining robust levels of Ser-linked modification even at HPF1 concentrations 10-to 20-fold lower than PARP1 (Fig. 2A, Fig. 1D). HPF1 was indeed reported to be approximately 20-fold less abundant than PARP1 in the cell ^16,22^, thus we considered that HPF1 could be tailored to biochemically operate at these sub-stoichiometric conditions. Even HPF1:PARP1 ratios of 1:50 and 1:100 yielded substantial amounts of Ser-linked modification. Similar results were also obtained for PARP2 (Fig. 2B), where PAR resistant to hydroxylamine treatment (i.e. Ser-linked) was observed even at the HPF1:PARP2 ratio of 1:100. However, the HPF1:PARP2 ratio of 1:1 was more efficient than the sub-stochiometric ratios at generating Ser-linked PAR. We first considered that HPF1 might impart a structural change on PARP1 that persists over time and thus does not require HPF1 to be continually engaged on each PARP1 molecule in order for Ser modification to occur. However, we were not able to isolate this putative, modified version of PARP1. Moreover, the recent structural analysis indicated that HPF1 directly contributes residue Glu284 to the PARP1/2 active site, thus indicating the requirement for a HPF1:PARP1/2 stoichiometry of 1:1 at the time of forming a Ser-linked ADP-ribose modification. We thus pursued a model in which HPF1 acts rapidly to modulate the output of multiple PARP1 molecules, moving from one molecule to the next to initiate Ser-linked modifications. In this model, HPF1 must act efficiently to engage consecutive PARP1 molecules, such that an excess pool of active PARP1 molecules is Ser-linked prior to the formation of Glu/Asp-linked modifications. Remarkably, we observed that HPF1 could efficiently modulate PARP1 output even when added at the same time as substrate NAD^+^ and at a HPF1:PARP1 ratio of 1:10 (Fig. 2C), indicating that HPF1 can rapidly engage PARP1, and also suggesting that initiation on Glu/Asp residues is a relatively slow process. We also considered the possibility that only a fraction of PARP1 molecules would be bound to DNA and active at any given time, as another potential explanation for efficient activity at the sub-stoichiometric HPF1:PARP1 ratios. However, HPF1 was also capable of regulating at a sub-stoichiometric ratio the constitutively active version of PARP1, PARP1ΔHD, which is active in the absence of DNA (Supp. Fig. 2). Though it is possible that a certain number of Glu/Asp modifications are present along with Ser modifications on PARP1 molecules when HPF1 is present at sub-stochiometric levels, the majority of the modifications would have to be on Ser to explain the hydroxylamine protection results.

**Figure 2.**
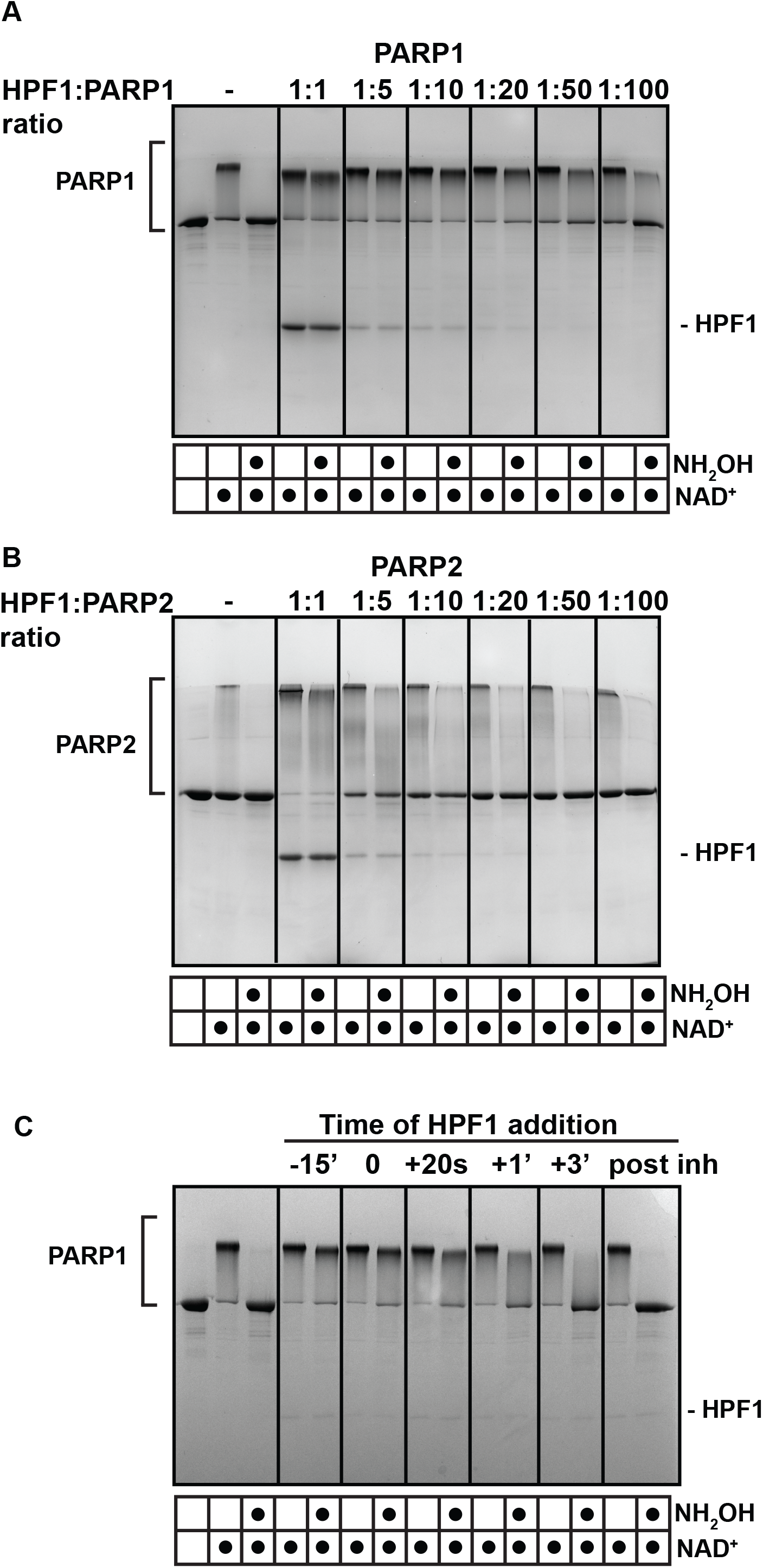
HPF1 works efficiently at sub-stochiometric ratios relative to PARP1 and PARP2. A) PARP1 (1 μM) or B) PARP2 (1 μM) was incubated with various amounts of HPF1 at the ratios indicated in the presence of DNA (1 μM) for 10 min at RT. 500 μM NAD^+^ was added for 5 min and reactions were quenched with 500 μM PARPi (olaparib or talazoparib). Where indicated, reactions were treated with hydroxylamine (NH_2_OH) for one hour. Reactions were resolved by SDS-PAGE and treated with Imperial Stain. C) PARP1 (1 μM) was mixed with HPF1 (1 μM) at various time points relative to NAD^+^ addition (500 μM). For the -15’ reaction, HPF1 was incubated with PARP1 for 15 min prior to NAD^+^ addition. For the 0 reaction, HPF1 was added at the same time as NAD^+^. For the + reactions, HPF1 was added after NAD^+^ at the time indicated (20 s, 1 min, 3 min). For “post inh”, HPF1 was added after the reaction was quenched with PARP inhibitor. Reactions were then treated as in panels A and B.

We evaluated HPF1 binding to PARP1 to gauge whether the kinetics of this interaction were consistent with the proposed model. Using SPR, we measured HPF1 interaction with PARP1 in complex with a biotinylated SSB DNA that was immobilized on a biosensor through streptavidin capture (Fig. 3). After PARP1 was allowed to associate with SSB DNA on the biosensor surface, HPF1 was injected during an early segment of the PARP1 dissociation phase, when PARP1 was still retained on the surface at high levels. HPF1 weakly interacted with the PARP1-DNA complex, but the interaction was greatly enhanced by the presence of the NAD^+^ mimic EB-47, which supports the HD conformation that exposes the ART for optimal HPF1 binding (Fig. 3A). These results are consistent with a recent qualitative analysis of the HPF1-PARP1 interaction over gel filtration in the presence of DNA and EB-47 ^18^. We observed a similar but less robust interaction when using the non-hydrolyzable NAD^+^ analog called BAD, which acts like EB-47 to influence HD conformation, but has a lower binding affinity for PARP1 (Fig. 3A). It is notable that after the HPF1 dissociation phase, PARP1 was retained at a higher level on the surface relative to PARP1 that underwent dissociation in the presence of buffer instead of HPF1. We interpreted this observation to indicate that HPF1 had further stabilized PARP1 on DNA and thus slowed the rate of dissociation. As the PARP1-EB-47 complex represents the active form of PARP1 and was the most stable species, we quantified HPF1 interaction with PARP1-DNA by performing SPR analysis with EB-47 in the system buffer. The presence of EB-47 also had the effect of stabilizing the baseline of PARP1 interaction with DNA during the dissociation phase. The steady state levels of HPF1 binding at different concentrations indicated a binding affinity of 550 nM (Fig. 3B, 3C). In comparison, the binding affinity of HPF1 for PARP1-DNA (no EB-47), or the ART domain alone (deleted HD), were reported as 3.5 µM and 1.5 µM, respectively ^23,19^. We also analyzed the kinetics of the interaction. HPF1 exhibited a rate of association (*k*_a_) of 1 × 10^5^ M^-1^second (s)^-1^ and a rate of dissociation (*k*_d_) of 0.06 s^-1^ (corresponding to K_D_ of 540 nM). The *k*_d_ of 0.06 s^-1^ corresponds to a half-life of roughly 12 s. We reversed the SPR setup and attached HPF1 to the biosensor using amine coupling, and observed similar results regarding the requirement of DNA and an NAD^+^ mimic for robust interaction, and we also verified that the H826E mutant abolished the HPF1-PARP1 interaction (Supp. Fig. 3). Overall, the SPR analysis indicated that HPF1 interaction with PARP1 is “hit and run” with rapid association and dissociation, rather than an extended dwell time with PARP1 in the activated state. The results are consistent with our model where one HPF1 molecule visits several PARP1 molecules and participates in the catalysis of Ser ADP-ribosylation before PARP1 performs modification of Glu/Asp residues.

**Figure 3.**
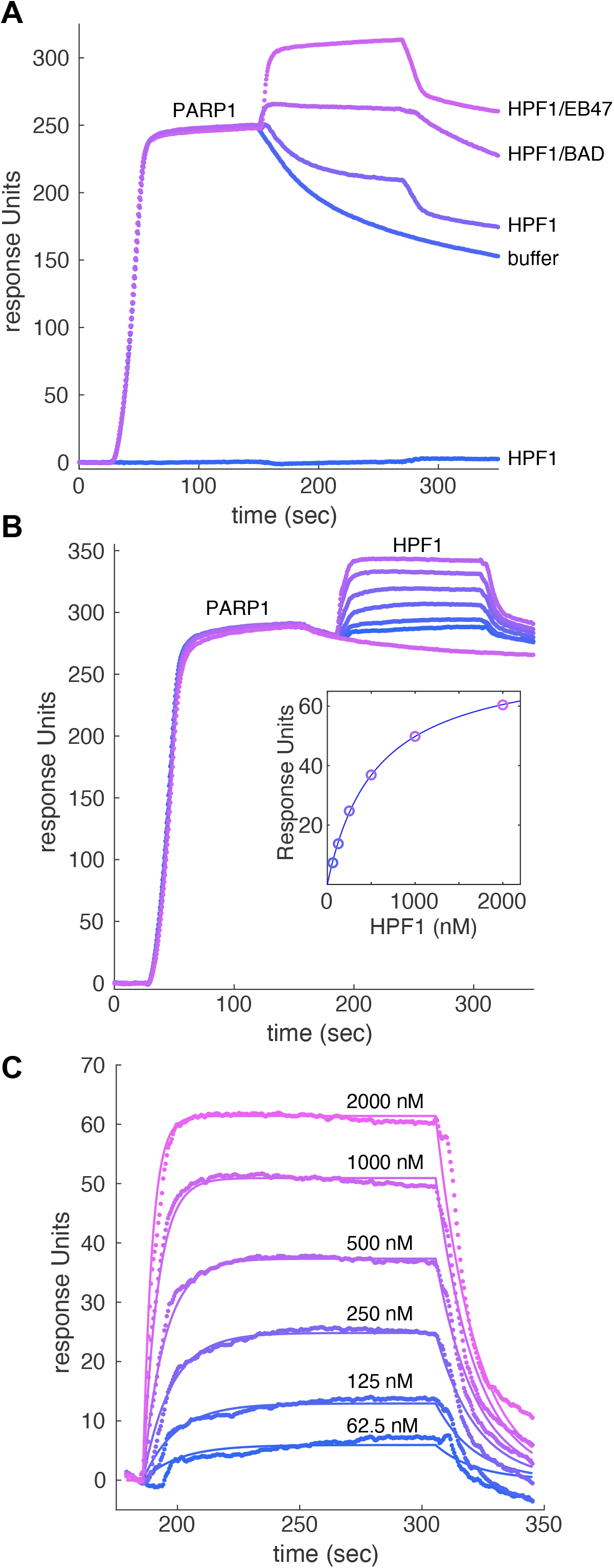
Kinetics of HPF1 interaction with the PARP1/DNA complex. A) A DNA SSB was immobilized on a biosensor chip using streptavidin/biotin capture. PARP1 (40 nM) was flowed over the chip until near saturation, thus forming a PARP1/DNA complex. The surface was then immediately exposed to the following solutions: buffer, HPF1, HPF1 with BAD, or HPF1 with EB-47. HPF1 alone did not interact with the DNA SSB surface. B) HPF1 was passed over the PARP1/DNA complex at the following concentrations: 0 (buffer only), 62.5, 125, 250, 500, 1000, and 2000 nM. In these experiments, the buffer was supplemented with 5 µM EB-47 to provide a more stable baseline during the PARP1 dissociation phase. The inset plots the plateau value achieved at each HPF1 concentration, and the line indicates the fit for a 1:1 binding model (K_D_ of 550 nM). C) The association and dissociation phases of the HPF1 injections from panel B were fit to a 1:1 binding model, yielding a rate of association (*k*_a_) of 1 × 10^5^ M^-1^second (s)^-1^ and a rate of dissociation (*k*_d_) of 0.06 s^-1^ (corresponding to K_D_ of 540 nM).

Our model also predicts that the HPF1-dependent Ser modification is faster than Glu/Asp modification. We observed indeed a stimulation of PARP2 initiation events in the presence of HPF1, visualized by a rapid decrease in intensity of the unmodified PARP2 band as early as 10 s in the presence of HPF1 (Fig. 4A). In contrast, a similar decrease in unmodified band intensity is only observed after 5 minutes (min) in the absence of HPF1 (Fig. 4A). In the case of PARP1, HPF1 stimulation of catalysis is not clearly observed when considering global activity on SDS-PAGE (Fig. 4B). In fact, we noted that HPF1 limits the length of polymer produced by PARP1, as previously described ^17^. HPF1 inhibition of PAR growth is more and more pronounced as the concentration of HPF1 was increased in PARP1 and PARP2 activity assays, indicating that HPF1 has the capacity to severely restrict elongation when used at elevated concentrations (Fig. 4B, C). This restriction is easily explained by the fact that HPF1 binding blocks access to the ADP-ribose acceptor site ^18,19^. To more directly focus on the HPF1 effect on PARP1 initiation reactions (as opposed to both initiation and elongation), we quantified PARP1 initiation rates in the presence and absence of HPF1. To accomplish this analysis, we performed ADP-ribosylation reactions for various time points and treated the quenched reactions with poly(ADP-ribose) glycohydrolase (PARG), an enzyme that digests the PAR polymer but leaves the initial ADP-ribose attached to PARP1 ^24^. The resulting mono(ADP-ribose) modifications on PARP1 were detected in a Western blot using a binding reagent that recognizes mono(ADP-ribose) ^25^. Strikingly, we observed in the presence of HPF1 that initiation was near saturation at 30 s, whereas initiation in the absence of HPF1 did not reach saturation until greater than 300 s (Fig. 5A, Supp. Fig. 4A). Using the early time-points in the linear region to estimate a rate of reaction, we observed that HPF1 increases the rate of initiation by roughly 50-fold relative to PARP1 alone (Supp. Fig. 5A). Moreover, the signal plateau reached by PARP1 in the presence of HPF1 is higher than the plateau reached in the absence of HPF1 (Fig. 5A, compare 300s and 600s with and without HPF1). Together, the data indicate that more initiation events take place in the presence of HPF1, and that the rate of accumulating initiation events is more efficient in the presence of HPF1.

**Figure 4.**
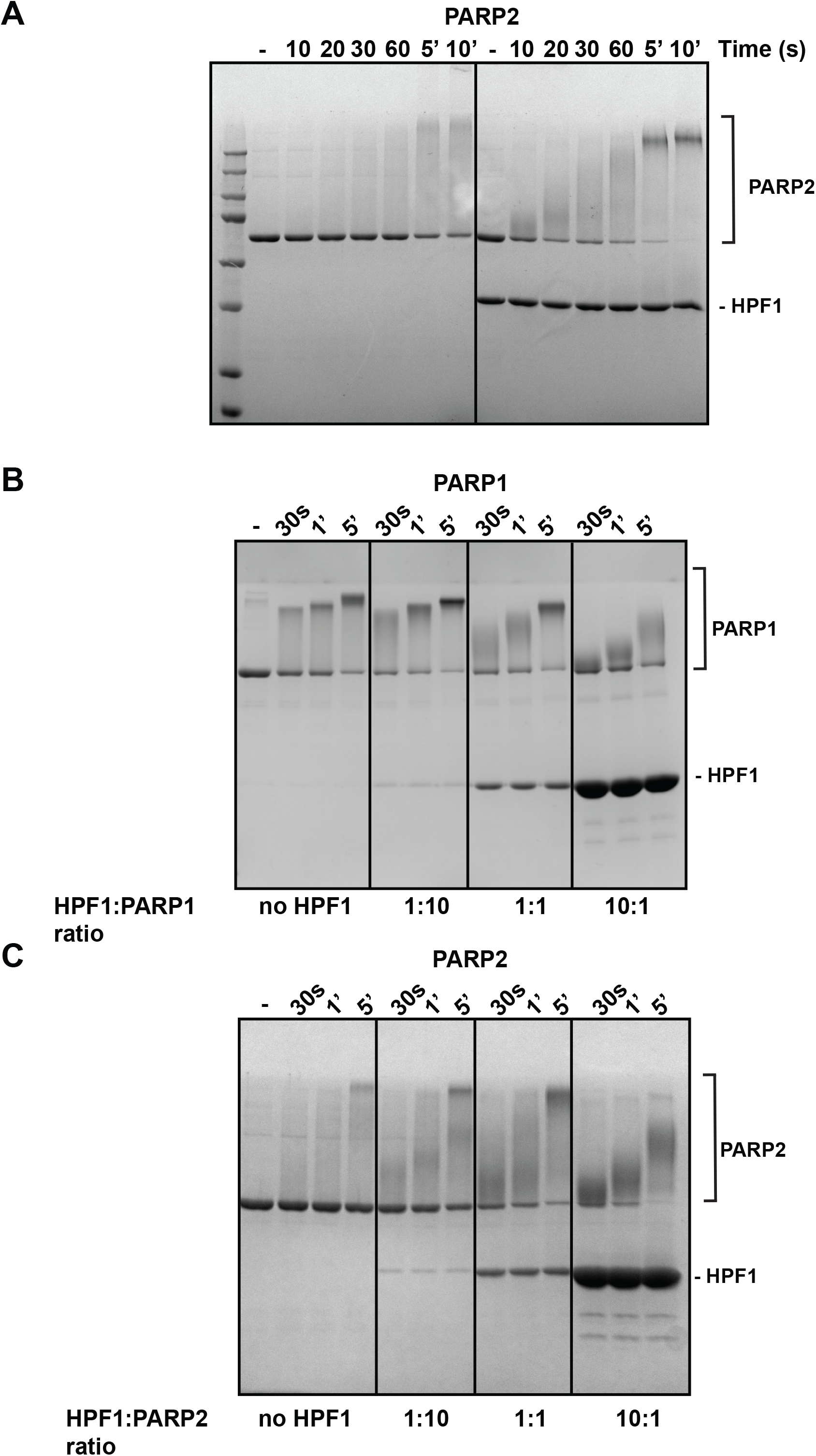
HPF1 restricts elongation and stimulates initiation in PARP2. A) PARP2 (1 μM) was incubated with HPF1 (1 μM) for 10 min at RT in the presence of DNA (1 μM), where indicated. 500 μM NAD^+^ was added for various time points (except in the “-” reaction), and reactions were quenched with SDS-PAGE loading buffer, resolved by SDS-PAGE, and treated with Imperial stain. B) PARP1 (1 μM) was incubated with HPF1 at various ratios as indicated in the presence of DNA (1 μM) for various time points with 500 μM NAD^+^. Reactions were processed as in panel A. C) Same as in panel B for PARP2.

**Figure 5.**
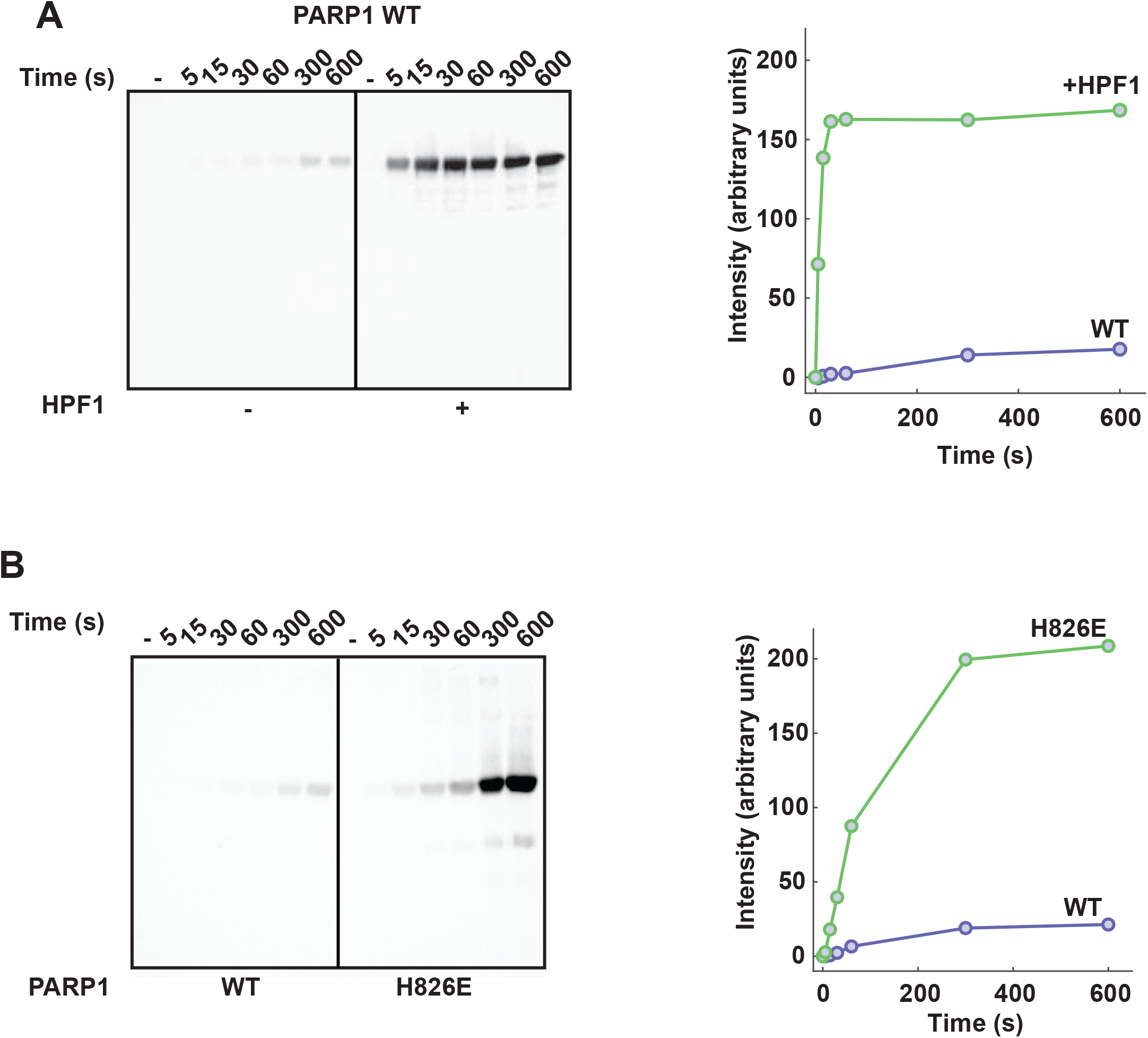
HPF1 stimulates initiation in PARP1. A) PARP1 (1 μM) was incubated without or with HPF1 (0.1 μM) for 10 min at RT in the presence of DNA (1 μM). 1 mM NAD^+^ was added for various time points as indicated, and reactions were quenched with 500 μM talazoparib. PARG (1 μM) was added and incubated for 1 hour at RT. Reactions were resolved by SDS-PAGE and a western blot was performed using a mono ADP-ribose binding reagent. The bands corresponding to mono ADP-ribosylated PARP1 were quantified using ImageJ and plotted as intensity over time. B) Reactions were performed as in panel without HPF1 and comparing PARP1 WT to PARP1 H826E.

We considered that the HPF1 block on the PARP1 elongation site could in effect divert the PARP1 active site to carry out more initiation events, thus explaining the observed increase in maximum signal. To test this reasoning, we used the PARP1 mutant H826E as a mimic of HPF1 disruption of the elongation site. As previously noted, H826E produces shorter polymers than WT due to the fact that His826 forms part of the ADP-ribose acceptor site (Fig. 1A, 1D). The rate of initiation for the H826E mutant is about 20-fold higher than WT PARP1 (Fig. 5B, Supp. Fig. 4B, Supp. Fig. 5B), and the maximum signal for this mutant was also higher than WT. The addition of HPF1 did not affect the activity of H826E (Supp. Fig. 6), consistent with the fact that HPF1 does not bind to this PARP1 mutant (Supp. Fig. 3)^18^. Though the maximum signal of H826E was higher than WT PARP1, it did not reach the level observed for WT PARP1 in the presence of HPF1 (Supp. Fig. 6). Together, these results suggest that restricting elongation indeed has the consequence of diverting PARP1 activity toward initiation events, and this mode of regulation can be invoked by HPF1 or through mutagenesis. However, the HPF1 effect on initiation is not due solely to restriction of elongation, since HPF1 substantially stimulates the rate at which PARP1 reaches saturation in our analysis of initiation sites.

HPF1 also modulates PARP1 catalytic output by increasing the extent of trans-modification of other factors, such as histones, relative to auto-modification ^17^. For example, histone H3 is not appreciably trans-modified by PARP1 unless HPF1 is present, and there is a substantial decrease in PARP1 auto-modification in the presence of HPF1 and histone H3 or histone octamer ^16,17^. To determine if HPF1 affects PARP1 trans-ADP-ribosylation at sub-stochiometric ratios at the initiation step, we incubated PARP1 with histone octamer in the presence and absence of HPF1, and treated the quenched reaction with PARG (Supp. Fig. 5C and Supp. Fig. 4C). Notably, the histone octamer inhibited PARP1 auto-modification even in the absence of HPF1. In the presence of HPF1 at a 1:10 ratio, PARP1 auto-modification was greatly reduced and one or more histones were modified indicating that even at sub-stochiometric ratio, HPF1 is able to modulate PARP1 catalytic output from auto-to trans-modification.

We undertook HXMS analysis to gain insights into HPF1 influence on PARP1 dynamics, in particular the dynamics of the HD domain and DNA binding regions. HXMS monitors the exchange of backbone amide hydrogens of a protein and thereby reports on protein structure and dynamics. Using this technique, we previously identified PARP1 regions that exhibit dramatic increases in HX, as well as regions that exhibit decreases in HX, when PARP1 is bound to DNA ^12^. The increases in HX locate to specific HD helices and relate to the allosteric mechanism that permits NAD^+^ binding, and the decreases in exchange locate to PARP1 regions that directly engage DNA or form domain-domain contacts. More recently, we used HXMS to study the effect of NAD^+^ analogs and PARP inhibitors on the dynamics of the PARP1 complex with DNA ^11,26^. Here, PARP1 was assembled on a DNA structure that models a DNA SSB, and the effect of adding HPF1 was analyzed through a full-time course HXMS experiment (from 10^1^ to 10^5^ s). Comparing HXMS data at 10 s of the PARP1/HPF1/DNA complex to the PARP1/DNA complex, we observed an increase in HX for the peptides in helix B (676-690) and a peptide at the C-terminus of helix F (771-778) of the HD (Fig. 6A-D and Supp. Fig. 7). These very same helices partially unfold when PARP1 is bound to a DNA break ^12^. Strikingly, differences in HX rates of the PARP1/DNA/HPF1 complex when compared to PARP1/DNA complex were only seen at 10 s and 100 s (see 100 s timepoint in Supp. Fig. 8). Our results indicate that HPF1 pushes the HD further toward the unfolded state, similar to the effect of NAD^+^ analogs.

**Figure 6.**
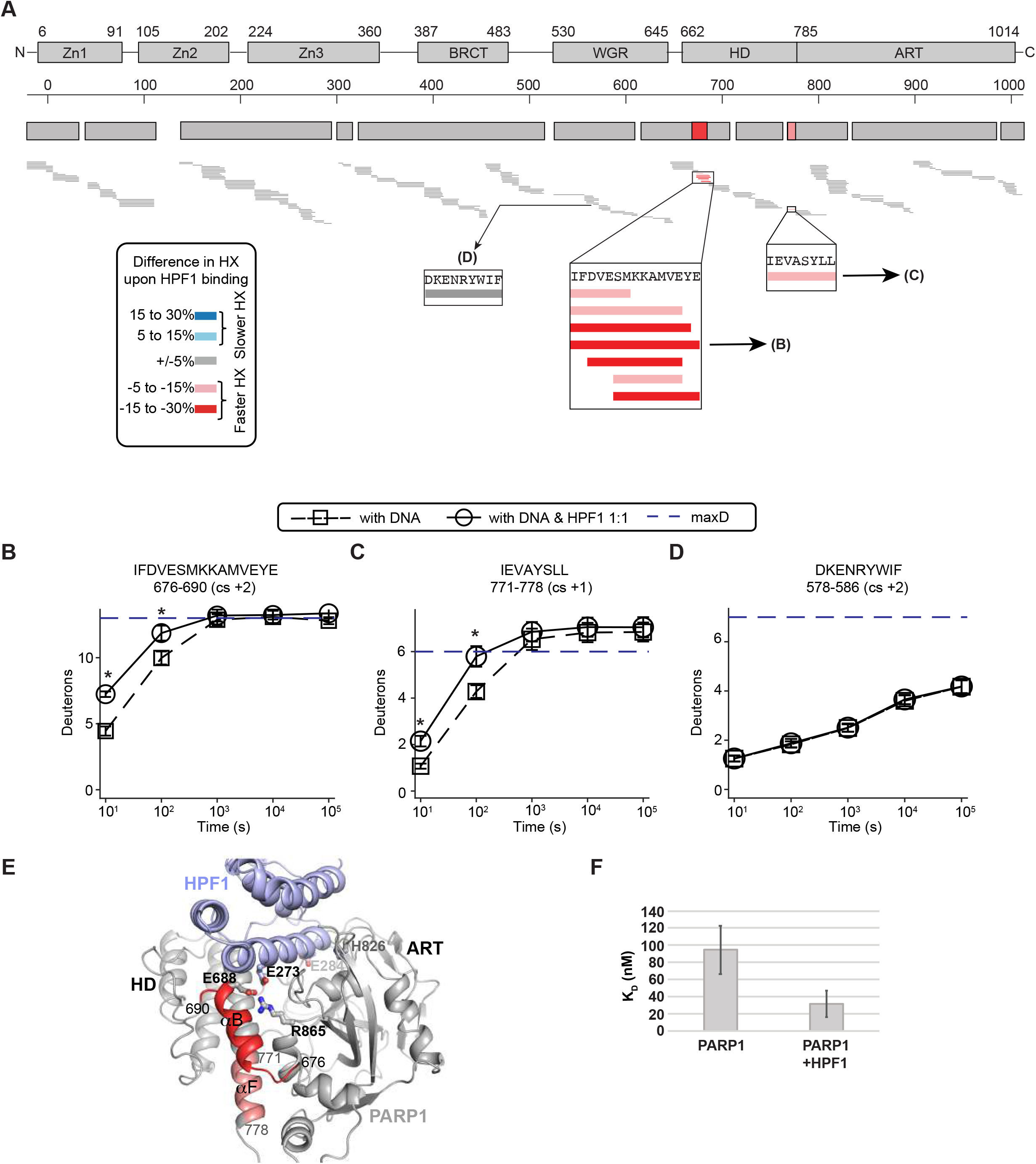
HPF1 further destabilizes the HD when PARP1 is bound to a DNA SSB. A) An HXMS difference plot obtained by subtracting the percent deuteration of the PARP1/DNA/HPF1 complex (at 1:1 ratio of HPF1:PARP1) from PARP-1/DNA complex at 10 s. Each horizontal bar represents a PARP1 peptide. Most of the peptides (grey) in the PARP1/DNA/ HPF1 complex have similar HX rates when compared to the PARP1/DNA complex. However, peptides in the αB and αF helices of HD showed faster exchange (red). The white regions in the difference plot represent gaps in the peptide coverage. B) HX of a representative peptide from the αB helix in panel A for PARP1/DNA and PARP1/DNA/HPF1. C) HX of a representative peptide from the αF helix in panel A for PARP1/DNA and PARP1/DNA/HPF1. D) HX of representative peptide from WGR domain in panel A for PARP1/DNA and PARP1/DNA/HPF1. Experiments were performed in triplicate. In panels B-D, the dotted lines indicate the number of deuterons measured with pre-denatured PARP1 (i.e. full exchange), and SD is represented by the error bars. In panels B-C, asterisks indicate *P* < 0.05 (*t*-test between PARP1/DNA and PARP1/DNA/HPF1 complexes). E) Consensus HXMS data from panel A mapped onto the structure of the catalytic domain of PARP1 (from 4dqy), which was modeled in complex with HPF1, based on the PARP1CATΔHD complex with HPF1 (6tx3). F) DNA binding affinity of PARP1 in the absence and presence of HPF1.

In contrast to NAD^+^ mimics, HPF1 did not induce an increase in protection in PARP1 regions directly involved in DNA binding. However, HPF1 did increase PARP1 affinity for DNA by about 3-fold in a fluorescence polarization DNA binding experiment (Fig. 6F). These results are consistent with the SPR results where the addition of HPF1 increased PARP1 retention on DNA (Fig. 3A). These data suggest that HPF1 destabilization of the HD does exert an allosteric effect on PARP1 binding to DNA, similar to the binding of Type I inhibitors such as EB-47 ^26^, the NAD^+^ analog BAD ^11^, and the veliparib derivative UKTT15 ^26^, albeit to a lesser extent. The structure of PARP1 CAT ΔHD bound to HPF1 shows that PARP1 residue Arg865 is repositioned to interact with HPF1 residue Glu273 when compared to the structure of PARP1 CAT without HPF1, in which Arg865 interacts with HD residue Glu688 located on helix B (Fig. 6C). Breaking the connection between the HD and the ART by HPF1 could promote HD conformational flexibility and thereby increase PARP1 retention on DNA. The findings of a transient interaction with PARP1 and a massive stimulation in the PAR-chain initiation step, along with the prior report that the HD must be removed to allow stable binding ^18^, suggest a model where the HD flexibility first permits HPF1 to recognize only PARP1 molecules engaged with a DNA break, but then stabilizes PARP1 on the break long enough (seconds timescale) to initiate PAR chains on local target serine residues. The idea of HPF1 stabilizing PARP1 on a DNA break initially seems at odds with the finding that genetic removal of HPF1 leads to PARP1 retention for several minutes on large lesions of DNA damage ^17^. The answer to this paradox may lie in a role for serine-linked PARylation in automodification-dependent release from these lesions that is defective in the absence of HPF1.

Structural insights into full-length PARP1 interaction with HPF1 are still lacking. Therefore, we used negative stain EM to provide first views of a PARP1-HPF1-DNA complex. PARP1 was incubated with a DNA SSB containing a one-nucleotide gap in the presence of EB-47, both with and without HPF1. The EM analysis indicates that PARP1 binds to SSB-DNA as a monomer in the absence and presence of HPF1 (Fig. 7). Additionally, even though the PARP1/DNA/HPF1/EB47 complexes were incubated at stochiometric ratios of proteins and DNA, only a fraction of the PARP1/DNA complexes observed on the grids (roughly 10%) contained HPF1. This result is consistent with our “hit and run” mechanism, where HPF1 assembles and disassembles rapidly from the PARP1/DNA complex. 2D class averages provided views in which certain PARP1 domains could be inferred based on knowledge from X-ray and NMR structures of PARP1 fragments, and the localized domains represent the most stable components of the complex: the Zn1, Zn3, WGR and CAT domains (Fig. 7). The 2D averages did not suggest a location for the BRCT domain, which we infer to remain flexibly tethered to the rest of the complex. Overall, the 2D classification indicated that although PARP1 forms a largely globular assembly, there is inherent flexibility within the assembly that prevents the construction of confident 3D maps. The DNA was not clearly visible in the images, likely due to the DNA size and multiple domains enveloping the DNA structure. Additional density was observed on the CAT domain in the presence of HPF1, consistent with our interpretation of PARP1 domains and their arrangement. Initial modelling suggested two main conformations for the PARP1-HPF1-DNA complex, one where HPF1 is flexed away from the domains assembled on DNA, and one where HPF1 approaches these domains (Fig. 7). In both conformations, the relative orientation between the CAT and HPF1 remained the same. The negative stain EM imaging thus indicates that HPF1 is incorporated into the PARP1 assembly of domains on a DNA break, with the potential to influence the overall stability of the PARP1-DNA complex. Moreover, HPF1 is positioned within the complex in a way that could influence PARP1 capacity to automodify, and therefore play a role in the regulation of *cis versus trans* modification.

**Figure 7.**
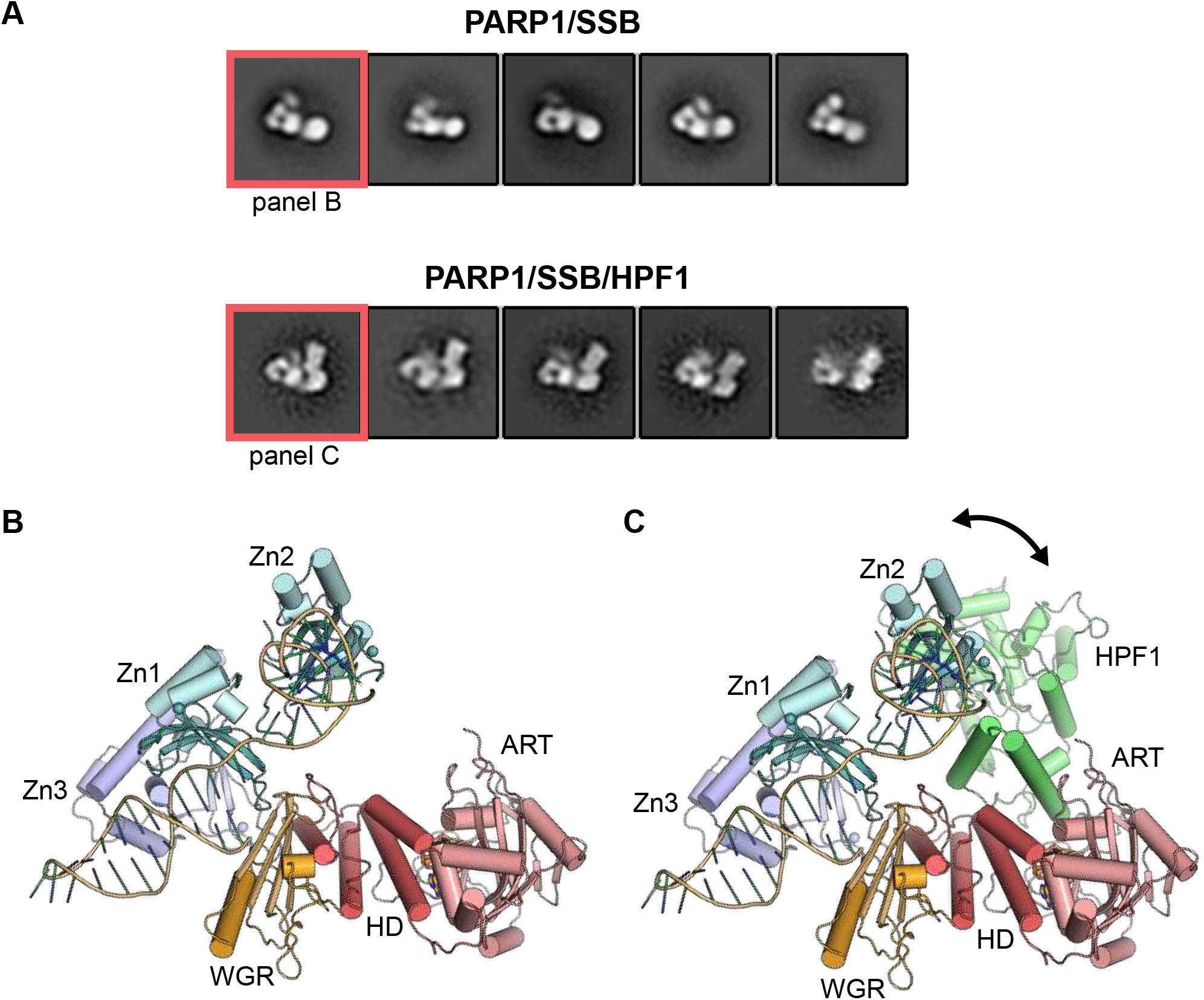
Architecture of the HPF1 complex with full-length PARP1 bound to a DNA break. The PARP1/DNA SSB complex was analyzed by negative stain electron microscopy in the absence and presence of HPF1. A) 2D class averages provided views of the complexes that could be interpreted with the aid of published structures of PARP1 domains (Zn1, Zn2, Zn3, WGR and CAT (HD/ART)) on DNA as shown in B), and with HPF1 in C). The 2D averages did not suggest a location for the BRCT domain, which we infer to remain flexibly tethered to the rest of the complex. Consistent with our interpretation of PARP1 domains, the addition of HPF1 adds to the globular end of the protein interpreted as the ART (panel C). The orientation of HPF1 with respect to the catalytic domain remains relatively fixed, whereas the orientation of HPF1 relative to the PARP1 regulatory domains appears to adopt two different conformations, highlighted by the arrow in panel C.

## DISCUSSION

The recent determination of high-resolution structures of PARP1 and PARP2 bound to HPF1 ^18,19^ have had a tremendous impact on our comprehension of HPF1 mode of action. HPF1 completes the active site of PARP1 by inserting catalytic residue Glu284, and this HPF1-PARP1 complex is essential for Ser ADP-ribosylation, the predominant type of modification in cells following DNA damage. It is thus tempting to imagine a stable and saturated HPF1-PARP1 complex to ensure that the appropriate target residues are modified. However, the strong inhibitory influence of HPF1 on the elongation reaction complicates this picture, and the cellular abundance of HPF1 does not match that of PARP1. The results presented in this study allow us to propose a new model for HPF1 regulation of PARP1/2 catalytic output (Fig. 8). Our data suggest that HPF1 is bound to PARP1 at the step of initiation, where it substantially stimulates the rate of attachment of the first ADP-ribose to Ser in comparison to the rate observed for the attachment of ADP-ribose to Glu/Asp in the absence of HPF1. For the time that HPF1 remains bound to PARP1, the elongation reaction is inhibited through a steric block on the acceptor site. Yet, the PARP1 active site remains available for additional initiation reactions on Ser residues, explaining why we observe a higher number of initiation events in the presence of HPF1. This shift in balance toward initiation events was also observed with PARP1 mutant H826E, which has a disrupted acceptor site and is restricted in PAR elongation. These results suggest that the elongation site could serve as a general engagement site for factors (proteins or small molecule ligands) to bind and modulate PARP1/2 catalytic output.

**Figure 8.**
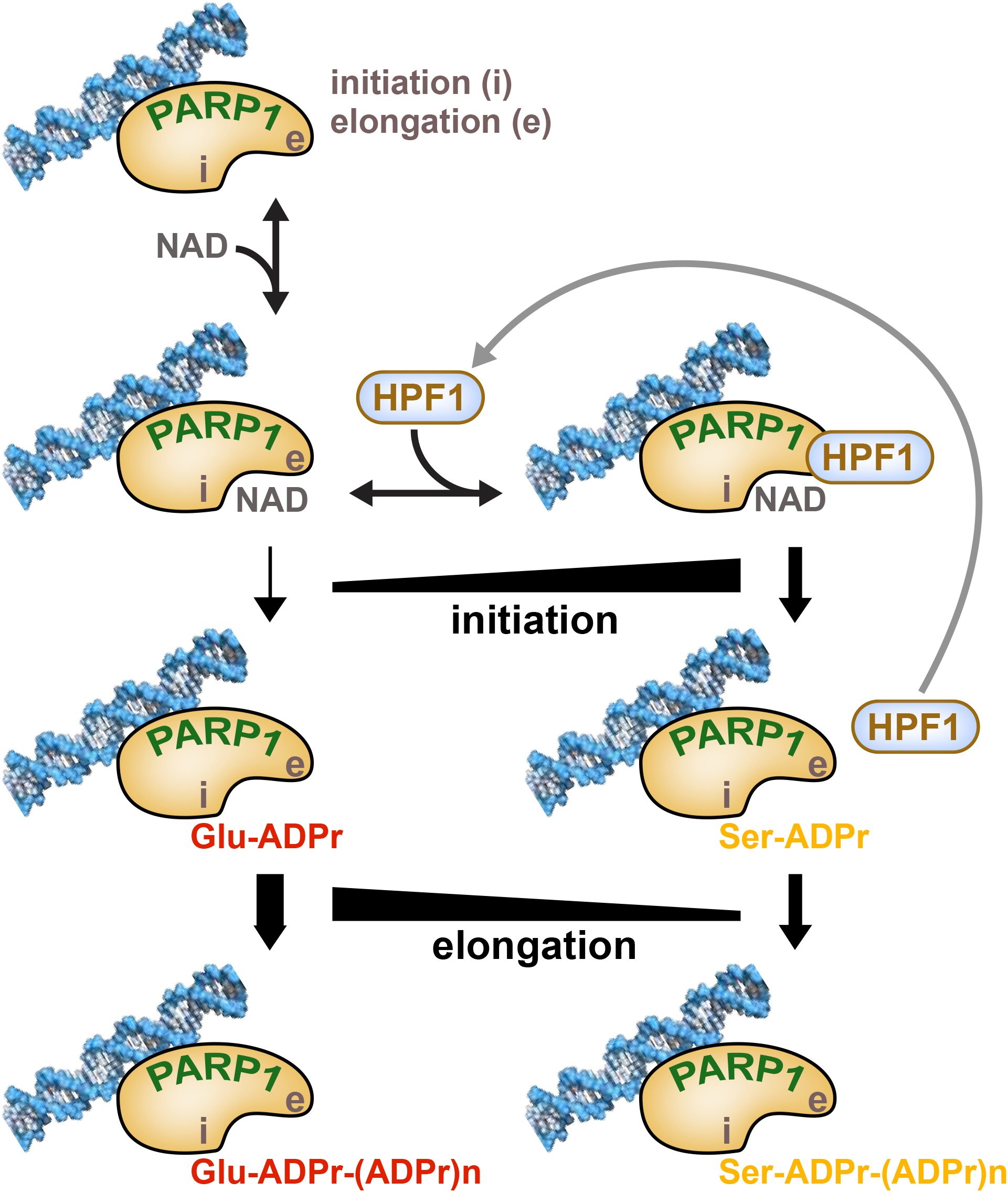
Model for HPF1 regulation of PARP1/2 catalytic output. HPF1 regulates the balance between initiation (i) and elongation (e) through a dynamic interaction with the catalytic domain and by accelerating the rate of initiation on Ser residues. Rapid HPF1 cycling between PARP1 molecules, indicated by the long arrow, allows the PARP1 population to be efficiently modified on Ser, and also prevents excessive inhibition of the elongation reaction to form PAR that is important for DNA damage signaling.

Our results showing that HPF1 functions efficiently at sub-stoichiometric ratios indicate that HPF1 must visit and initiate Ser modification on multiple PARP1 molecules before PARylation is initiated on Glu/Asp residues, hence the “hit and run” mechanism that we propose. Our SPR data show that indeed HPF1 can associate/dissociate rapidly from PARP1. Therefore, we propose that there is a fine balance between HPF1 remaining bound to PARP1 and blocking PAR chain elongation, and HPF1 dissociating and binding to a different PARP1 molecule to ensure that ADP-ribosylation takes place on Ser residues. It is notable that stochiometric amounts of HPF1, or an excess of HPF1, has the effect of strongly inhibiting global PARP1 and PARP2 PARylation activity (Fig. 5B), due to the steric blockage of the elongation/acceptor site, reducing more and more the size of the polymer as the concentration of HPF1 is increased. These results highlight that saturating the HPF1 interaction with PARP1 reduces catalytic output to a level that would be not be productive. The ratio of HPF1 to PARP1 in the cell could reflect a balance of ensuring efficient initiation of Ser modification, while also limiting the impact on ADP-ribose polymer production that is key to multiple aspects of DNA damage repair signaling. Our model also applies to trans-modification of histones; even at the HPF1:PARP1 ratio of 1:10, HPF1 supports the switch from auto-modification of PARP1 to trans-modification of histones as observed with the large decrease in signal observed on PARP1 in the presence of HPF1 and the histone octamer (Fig. 5D).

PARP1 initiation in the absence of HPF1 is rather inefficient (Fig. 5A). This observation is not readily apparent when observing the global activity on SDS-PAGE (*e*.*g*. Fig. 1C and Fig. 4B). Previous PARP1 studies without HPF1 have shown that the elongation step of PAR formation is at least 30-fold more efficient than the initiation step ^27–29^, and PAR size analysis from PARP1 reactions has indicated a distribution skewed toward long polymer sizes ^30^, which supports slow initiation but rapid elongation events. Thus, in the absence of HPF1, initiation on Glu is a relatively slow and inefficient event, but once a Glu initiation event has occurred, the elongation step is much more rapid, creating long chains of PAR that shift the PARP1 molecule on SDS-PAGE. Therefore, globally, the PARylation reaction appears to be quite efficient in the absence of HPF1. In the presence HPF1, the balance of initiation *versus* elongation is shifted and the rate of initiation is substantially increased, such that the complete pool of PARP1 molecules is initiated within seconds. This effect can be clearly observed in our analysis of initiation events (Fig. 5), and it can also be appreciated by observing the loss of un-modified PARP1 in the SDS-PAGE reactions at early time points (Fig. 4B). HPF1 thus elevates the rate of initiation substantially such that initiation is more in balance with elongation.

Stochiometric amounts of HPF1, or an excess of HPF1, has the effect of strongly inhibiting global PARP1 and PARP2 PARylation activity (Figs. 4B and 4C), due to the steric blockage of the elongation/acceptor site, reducing more and more the size of the polymer as the concentration of HPF1 is increased. HPF1 inhibition of elongation is likely to contribute to the shift toward initiation. The compact conformation of PARP1 on a DNA break was proposed to position residues for initiation of the ADP-ribose modification ^10,31^. Ser residues and Glu residues identified consistently in mass spectrometry analysis of automodification sites ^16,32–34^ are located in a region predicted to position near the active site of PARP1 ^10,31^. The Ser residues could be in the “most optimal” position for modification compared to Glu residues, and HPF1 simply allows these Ser residues to be utilized by completing the active site ^18^. However, the automodified region represents a lengthy linker region that is likely to be quite flexible, thus it is challenging to envision a rigidly fixed, “most optimal” region to be modified. The HPF1-PARP2 crystal structure identified a cleft near the active site termed the “canyon” that was proposed to specifically engage peptides to be modified. The peptide canyon could contribute to the acceleration of initiation events by providing a specific site of engagement for target residues, and a Lys-Ser motif is emerging as a signature sequence that could be engaged.

Overall, our study adds a new level of understanding of HPF1, and how it dynamically influences the initiation and elongation steps of PARP1/2 PARylation activity. These special properties allow HPF1 to regulate the balance of PARP1/2 catalytic output to match the requirements for ADP-ribose production in the cellular response to DNA damage, both in terms of the speed of catalyzing ADP-ribose modification and the size distribution of polymer formed. We also present HXMS and negative stain EM analysis that provide first insights into the structure and dynamics of full-length PARP1 in complex with HPF1, providing a framework for better understanding the special collaboration between these proteins in the DNA damage response.

## EXPERIMENTAL PROCEDURES

### Expression constructs and mutagenesis

PARP1 (residues 1 to 1014) and PARP2 (isoform 2, residues 1 to 570) were expressed from a pET28 vector with an N-terminal hexahistidine tag. PARP3 FL (isoform b, residues 1 to 533) was expressed from a pDEST17 vector with an N-terminal hexahistidine tag (gift from Dr. Ivan Ahel). The human HPF1 gene was synthesized for expression from a pET28 vector with an N-terminal SMT sumo-like tag. Human histones genes were synthesized for co-expression: H2A/H2B in vector pCDF Duet, and H3/H4 in vector pET Duet-1. Site-directed mutagenesis was performed using the QuikChange protocol (Stratagene) and verified by automated Sanger sequencing. PARP1ΔHD was generated as described in ^12^.

### Protein Expression and Purification

PARP1 WT and mutant proteins were expressed and purified as described previously ^8,12,35,36^ using Ni^2+^-affinity, heparin, and gel filtration chromatography. Purification of PARP2 and PARP3 was performed as described ^13^ using Ni^2+^-affinity, heparin, and gel filtration chromatography. Purification and reconstitution of PARP1ΔHD was performed as described ^12^ using the transpeptidase Sortase A. Histones H2A and H2B were co-expressed in *E. coli* and purified as a soluble dimer using anion exchange, heparin, and gel filtration chromatography. Histones H3 and H4 were co-expressed in *E. coli* and purified as a soluble tetramer using anion exchange, heparin, and gel filtration chromatography. The histone octamer was formed by mixing H2A/H2B with H3/H4 and further purification was performed on a Sephacryl S200 gel filtration column.

### SDS-PAGE PARP-1 Activity Assay

The SDS-PAGE activity assay was performed as described ^36^ using 1 μM PARP1 or PARP2, various amounts of HPF1 as indicated, 1 μM DNA (dumbbell with a central nick for PARP1, 28 bp duplex DNA with a 5’ Phosphate for PARP2), and 500 μM NAD^+^. Reactions were incubated for 5 min or as indicated before stopping the reaction with PARP inhibitor veliparib at 500 μM. Where indicated, reactions were treated with 1M hydroxylamine (NH_2_OH) for 1 hour at room temperature, then quenched with 0.3% HCl. SDS-PAGE loading buffer was added to the reactions prior to resolution on a 12% SDS-PAGE, which was treated with Imperial stain for visualization.

### Binding analysis using SPR

SPR was performed on a Reichert 4SPR biosensor equipped with a supplementary valve that allows for sequential injections. The system buffer was 25 mM HEPES pH 7.4, 250 mM NaCl, 0.1 mM TCEP, 1 mM EDTA, and 0.05% Tween20. The buffer was supplemented with 5 µM EB-47 for certain experiments as indicated. Streptavidin-coated chips (Reichert) were used to capture a DNA SSB bearing a biotin group (20 nM). PARP1 was flowed over the biosensor at 40 nM, and HPF1 was then injected over a range of concentrations in the absence or presence of EB-47 (5 µM) or BAD (300 µM). CM5 chips (Reichert) were used to immobilize HPF1 using standard amine coupling procedures. PARP1 (800 nM) was flowed over the HPF1-coated biosensor in the absence or presence of DNA (800 nM) and EB-47 (5 µM). All data was double-referenced to buffer and a control channel that did not contain the immobilized binding partner. The HPF1 titration onto PARP1/DNA SSB was processed and fit with a 1:1 binding model using TraceDrawer (Reichert).

### Western blot assay

PARP1 WT (1 μM) was incubated with DNA (1 μM; dumbbell with a central nick) without or with HPF1 (0.1 μM) and histone octamer where indicated (1 μM) at RT for 10 min as described ^36^. Reactions were started by adding NAD^+^ (1 mM) for various time points, stopped with PARP inhibitor talazoparib (500 μM) then treated with PARG (1 μM) for one hour at RT. SDS-PAGE (7.5% or 12 %) were loaded for experiments without histone octamer. When the experiments contained histone octamer, a 4-20% gradient SDS-PAGE was used to allow both PARP1 and histones to be detected on the same gel. The gels were transferred to a nitrocellulose membrane and the membrane was then blocked with 1% milk. The membrane was incubated with a mono ADP-ribose binding reagent (MABE 1076, Millipore Sigma, 1:2500) and next treated with a secondary antibody (goat anti-rabbit conjugated to HRP, Santa Cruz, 1:7000). The signal was revealed using ECL (Bio-Rad).

### HXMS

Prior to deuterium on-exchange, 2.6 μM of PARP1 was incubated for 30 mins with 5 μM of SSB DNA ^9^. For the PARP1/DNA/HPF1 complex, 2.6 μM HPF1 was added to the mixture and incubated for another 30 mins. Deuterium on-exchange was performed at room temperature by adding 5 μL of mixture to 15 μL of deuterium on-exchange buffer (10 mM HEPES, pD 7.0, 150 mM NaCl, in D_2_O, pD = pH + 0.4138) to yield a final D_2_O concentration of 75%. At specified time points, 20 μL aliquots were added into 30 μL of ice-cold quench buffer (1.66 M guanidine hydrochloride, 10 % glycerol, and 0.8 % formic acid to make a final pH of 2.4-2.5) and immediately frozen in liquid nitrogen until further use. Non-deuterated (ND) samples of PARP1 were prepared in 10 mM HEPES, pH 7.0, 150 mM NaCl buffer and each 20 μL aliquot was quenched into 30 μL of quench buffer. To mimic the on-exchange experiment, the fully deuterated (FD) samples were prepared in 75 % deuterium, but denatured under acidic conditions (0.5 % formic acid). The samples were incubated for 48 hours to ensure every amide proton along the entire polypeptide chain to undergo full exchange. 50 μL of samples were melted at 0 °C, rapidly injected into pepsin column and simultaneously pumped at initial flow rate of 50 μl min^-1^ for 2 min followed by 150 μl min^-1^ for another 2 min. Pepsin (Sigma) was coupled to POROS 20 AL support (Applied Biosystems) and the immobilized pepsin was packed into a 64 μL column (2 mm × 2 cm, Upchurch). The pepsin-digested peptides were trapped onto a TARGA C8 5 μm Piccolo HPLC column (1.0 × 5.0 mm, Higgins Analytical) and eluted through an analytical C18 HPLC column (0.3 × 75 mm, Agilent) with a 12-100% buffer B gradient at 6 μL/ min (Buffer A: 0. 1% formic acid; Buffer B: 0. 1% formic acid, 99.9% acetonitrile). The effluent was electrosprayed into the Exactive Plus EMR-Orbitrap (Thermo Fisher Scientific).

### HXMS analysis and plotting

To identify the likely sequence of parent peptides via the SEQUEST (Bioworks v3.3.1), the ND samples were injected into LTQ orbitrap XL, (Thermo Fisher Scientific) for tandem MS. A MATALAB based program, ExMS was used to identify the parental non-deuterated peptides through peptide envelope centroid values and chromatographic elution time ranges. HDExaminer software (v 2.5.0) was used next, which now uses the peptide pool information generated through ExMS to identify the deuterated peptides for every sample in the full-time course HXMS experiment. Each individual deuterated peptide is corrected for loss of deuterium label (back exchange after quench) during HXMS data collection by normalizing to the maximal deuteration level of that peptide in the fully deuterated samples. The median extent of back-exchange in our experiment is 26 %. The difference plot for the deuteration levels between any two samples was obtained through an in-house script written in MATLAB. The script compares the deuteration levels between two samples (e.g., PARP1/DNA complex and PARP1/DNA/HPF1 complex) and plot the percent difference of each peptide, by subtracting the percent deuteration of PARP1/DNA/HPF1 complex from PARP-1/DNA complex and plot according to the color legend in stepwise increments (as in Fig. 6A).

### Negative stain EM

A PELCO easiGlow (Ted Pella Inc.) was used to glow discharge carbon-coated copper grids (Electron Microscopy Sciences) prior to sample adsorption. Purified PARP1 was assembled with SSB DNA and EB-47 with or without HPF1 at 5 μM in 25 mM Hepes pH 8.0, 150 mM NaCl, 0.1 mM TCEP buffer. PARP1-DNA-EB-47 (5 μM) and PARP1-DNA-EB-47-HPF1 (2.5 μM) were adsorbed and stained with 1.5% uranyl formate using the “rapid flush” technique ^37^. Grids were imaged at 67,000x magnification with a FEI Tecnai 12 transmission electron microscope operated at 120 keV using a LaB6 filament and FEI Eagle 4k CCD (Electron Imaging Facility, Faculty of Dental Medicine, Université de Montréal). A total of 158 (complex lacking HPF1) and 165 (complex with HPF1) micrographs were collected using the SerialEM program ^38^. The micrographs were processed with RELION 3.1 ^39^ without contrast transfer function correction. Reference-free particle picking generated 203,168 (complex without HPF1) and 167,267 (complex with HPF1) particle coordinates. Pick locations were extracted in 80 pixel boxes and downsampled to 3.3 Å/pixel, corresponding to 264 Å box lengths. Iterative 2D classifications using 180 Å circular masks obtained a final dataset of 48,250 (without HPF1) and 25,335 (with HPF1) particles.

## ACKNOWLEDGEMENTS

We acknowledge support from the Canadian Institutes of Health Research (BMA436870 to J.M.P), the Natural Sciences and Engineering Research Council of Canada (RTI-2018-00894 to J.M.P), and the National Institutes of Health (R35GM130302 B.E.B). Efforts to characterize eukaryotic pathways relevant to human cancers are supported in part by National Cancer Institute grant Structural Biology of DNA Repair (SBDR; CA92584).

