## Supplemental Figures 1 to 8 for "HPF1 dynamically controls the PARP1/2 balance between initiating and elongating ADP- ribose modifications"

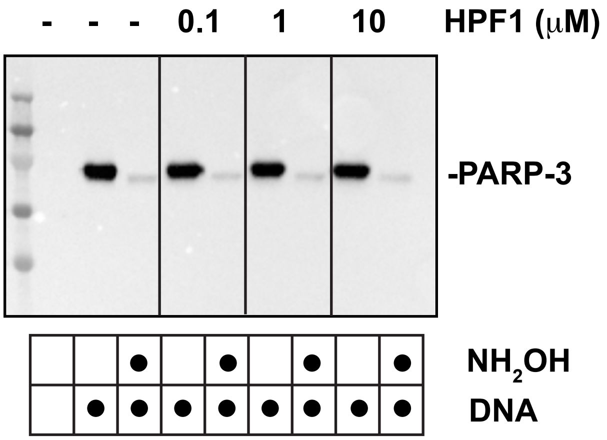


**Supplement Figure 1: HPF1 does not have an effect on PARP3 activity.** PARP3 (1 μM) was incubated with a dumbbell DNA containing a central nick (1 μM) with a 5' Phosphate group (5'P) with or without HPF1 at various concentrations for 10 minutes at room temperature (RT). 500 μM $NAD^+$ was added for 15 minutes and reactions were quenched with 10 μM PARP inhibitor (veliparib). Where indicated, reactions were treated with 1M hydroxylamine ($NH_2OH$) for one hour. Reactions were run on SDS-PAGE and a western blot was performed using a pan ADP-ribose binding reagent.

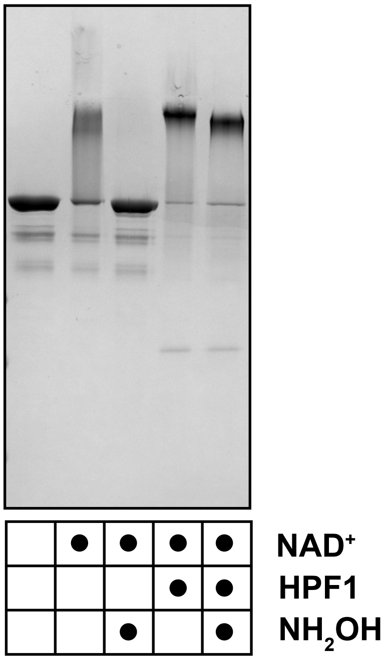


**Supplement Figure 2: HPF1 works at sub-stochiometric amount with the constitutively active PARP1 ΔHD in the absence of DNA.** PARP1 ΔHD (1 μM) was incubated with or without HPF1 (0.1 μM) for 10 minutes at room temperature (RT). 500 μM $NAD^+$ was added for 5 minutes and reactions were quenched with 500 μM PARP inhibitor (talazoparib). Where indicated, reactions were treated with 1M hydroxylamine ($NH_2OH$) for one hour. Reactions were run on an SDS-PAGE and stained with Imperial Stain.

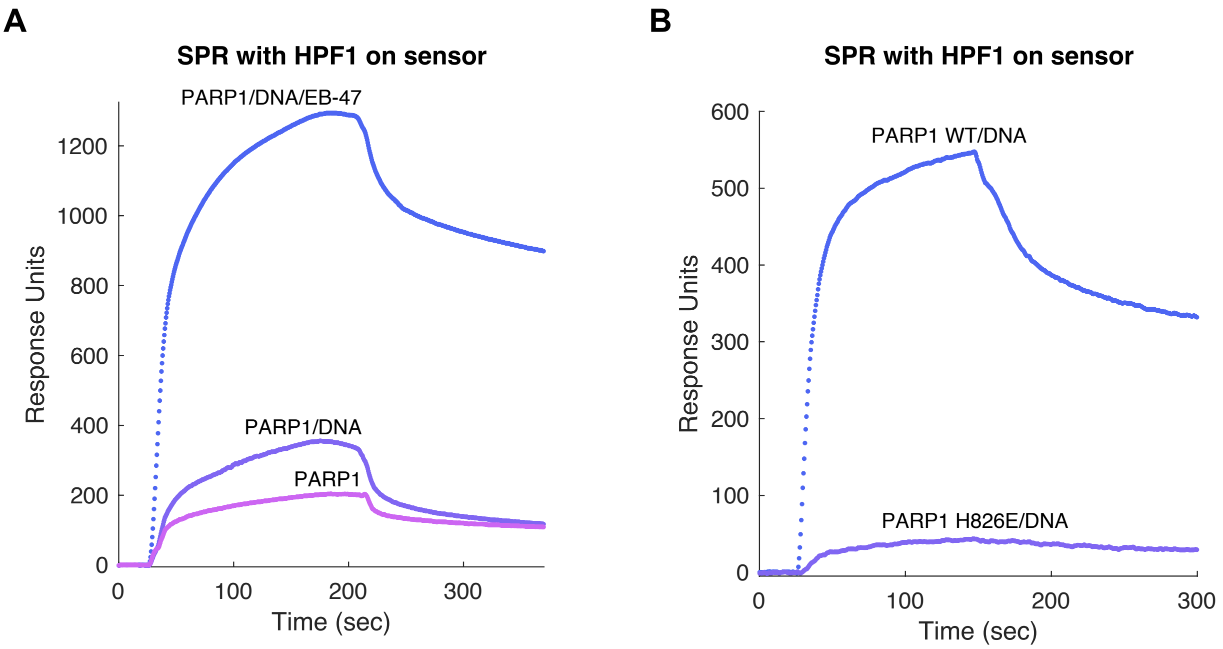


**Supplement Figure 3: HPF1 binds better to a PARP1/DNA complex in the presence of EB-47.** A) HPF1 was immobilized on a biosensor chip by amine coupling. PARP1 (800 nM) was flowed over the HPF1-coated biosensor in the absence or presence of DNA (800 nM) and EB-47 (5 μM). B) In a separate experiment, PARP1 WT or H826E mutant (1 μM) in the presence of DNA (1 μM) were flowed over an HPF1 chip with EB-47 (5 μM) in the system buffer.

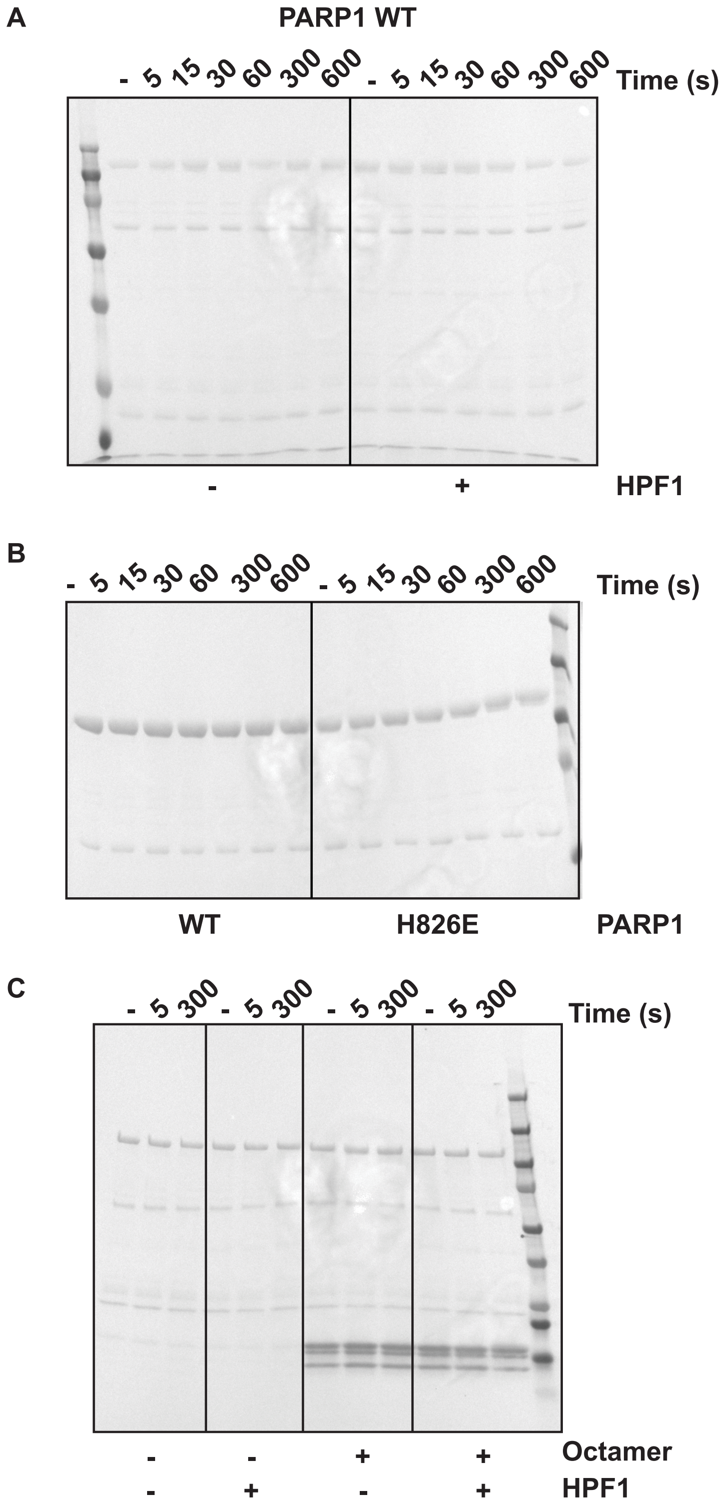


**Supplement Figure 4: Western Blots loading controls.** Ponceau-stained membranes corresponding to Western Blots shown in Figure 5 A, B and C.

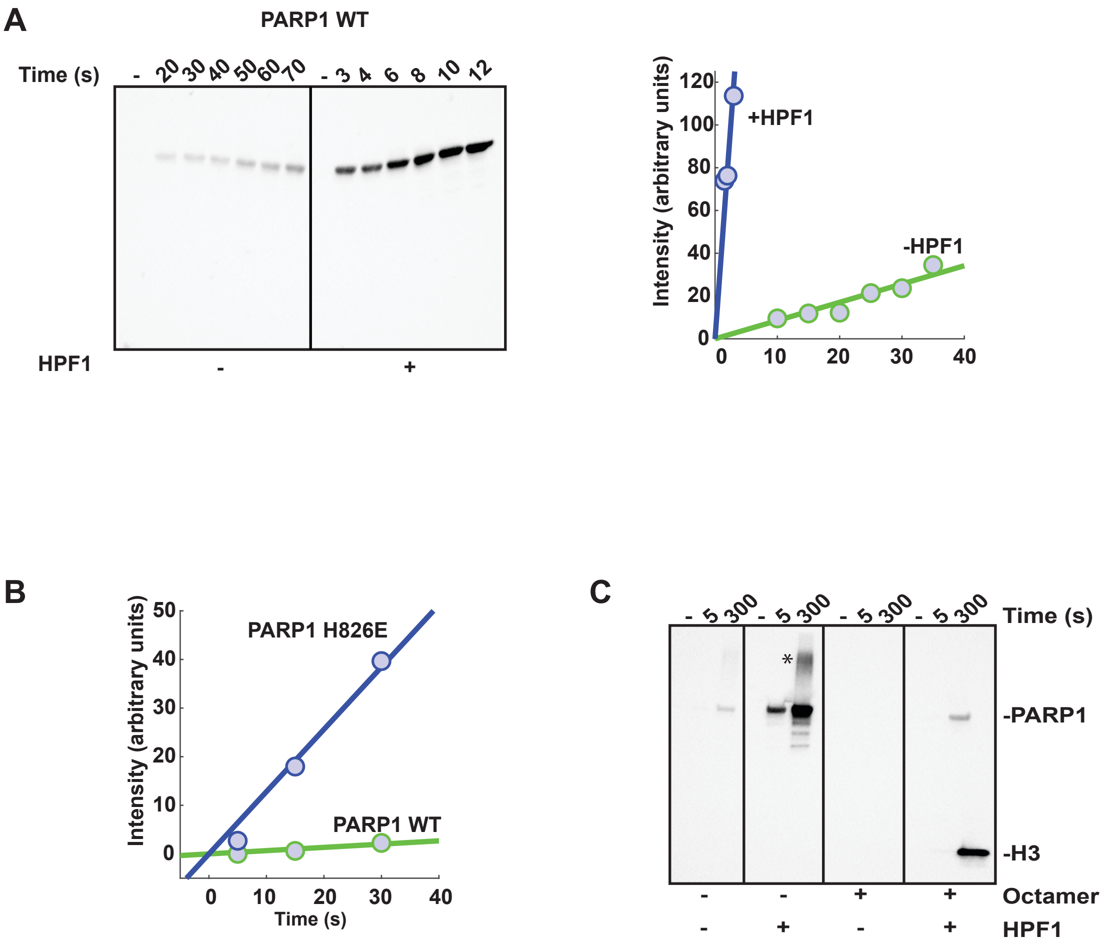


**Supplement Figure 5: HPF1 stimulates initiation by PARP1.** A) PARP1 (1 µM) was incubated without or with HPF1 (0.1 µM) for 10 minutes at RT in the presence of DNA (1 µM). 1 mM $NAD^+$ was added for various time points as indicated and reactions were quenched with 500 µM Talazoparib. PARG (1 µM) was added and incubated for 1 hour at RT. Reactions were run on SDS-PAGE and a western blot was performed using a mono ADP-ribose binding reagent. The bands corresponding to mono ADP-ribosylated PARP1 were quantified using ImageJ. Early time-points in the linear region were used to estimate a rate of reaction. B) Early time-points in the linear region from the experiment shown in the main Figure 5B were used to estimate a rate of reaction for PARP1 versus H826E. C) Reactions were performed as in A) but in the presence of histone octamer, where indicated. These reactions were run on 4 to 20% gradient gels and the additional band (denoted *) likely corresponds to very large, undigested PAR molecules that are usually not observed when using 12% gels such as in panels A and B, where they would not enter the resolving gel.

A

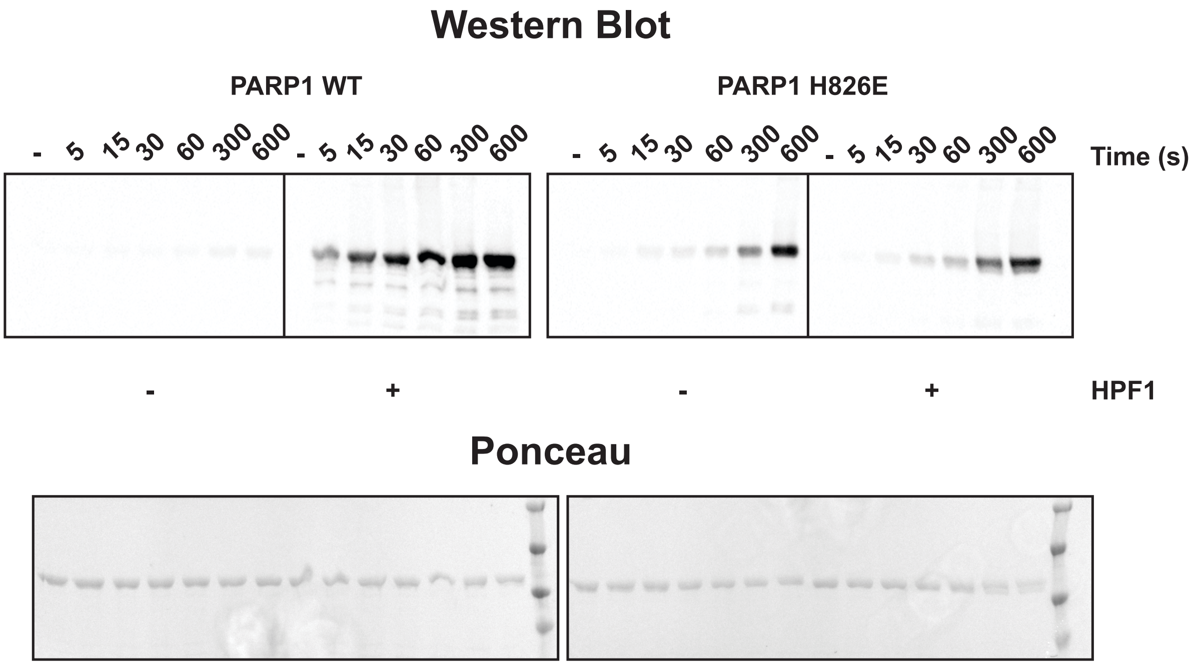


B

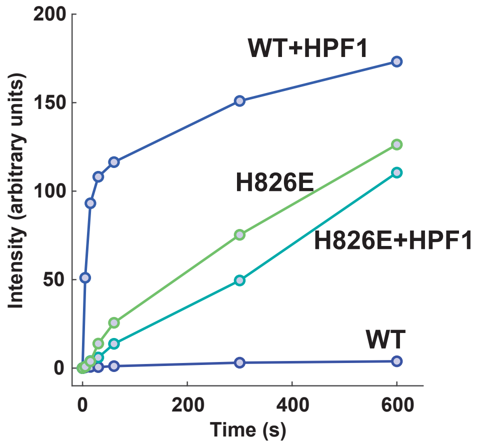


**Supplement Figure 6: HPF1 does not stimulate PARP1 mutant H826E.** A) PARP1 WT or H826E mutant (1 μM) was incubated without or with HPF1 (0.1 μM) for 10 minutes at RT in the presence of DNA (1 μM). 1 mM $NAD^+$ was added for various time points and reactions were quenched with 500 μM Talazoparib. PARG (1 μM) was added and incubated for 1 hour at RT. Reactions were run on SDS-PAGE the gels were cut so they could be transferred on the same membrane so that the signals could be compared and a western blot was performed using a mono ADP-ribose binding reagent. B) The bands corresponding to mono ADP-ribosylated PARP1 were quantified using ImageJ and the intensities were plotted over time.

**A**

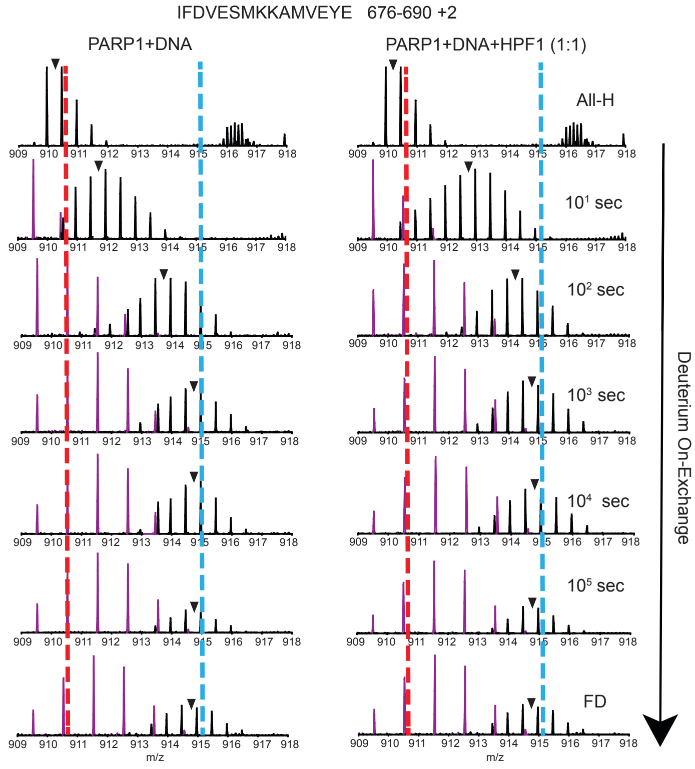
IFDVESMKKAMVEYE 676-690 +2
PARP1+DNA
PARP1+DNA+HPF1 (1:1)
All-H
10¹ sec
10² sec
10³ sec
10⁴ sec
10⁵ sec
FD
909 910 911 912 913 914 915 916 917 918
m/z
Deuterium On-Exchange

**B**

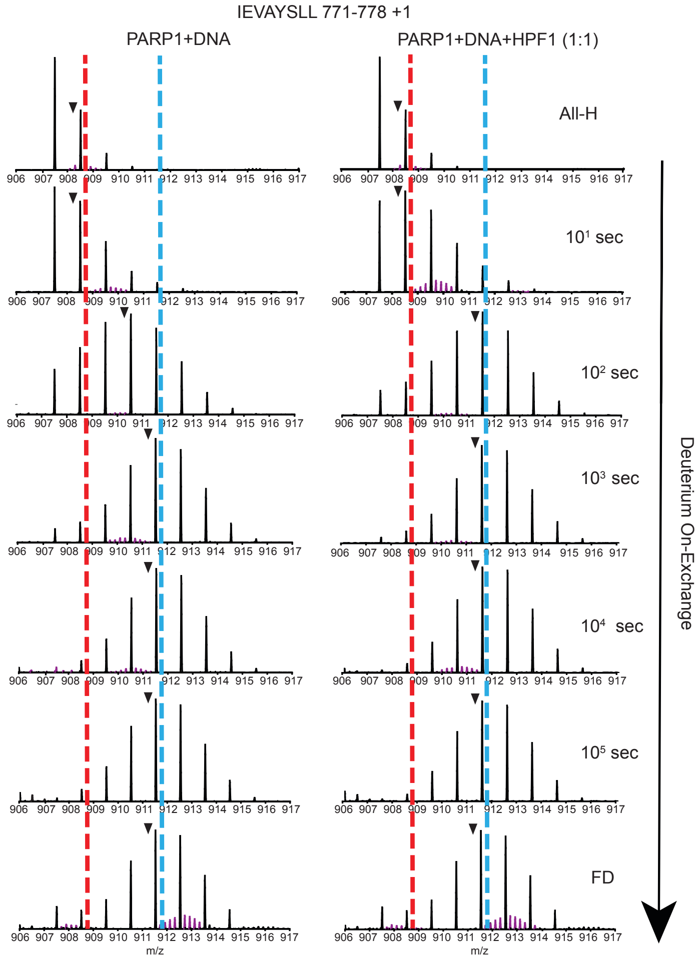
IEVAYSLL 771-778 +1
PARP1+DNA
PARP1+DNA+HPF1 (1:1)
All-H
10¹ sec
10² sec
10³ sec
10⁴ sec
10⁵ sec
FD
906 907 908 909 910 911 912 913 914 915 916 917
m/z
Deuterium On-Exchange

**Supplement Figure 7: Raw MS data of representative peptide from αB and αF helices of HD.** Spectra for PARP1/DNA complex and PARP1/DNA/HPF1 complex at 1:1 HPF1:PARP1 for the full-time course HXMS experiment is shown. All- H represents the non-deuterated sample. FD represents the fully- deuterated sample. Black isotopic envelopes represent the αB peptide (A) and αF helices (B) of HD, whereas purple isotopic envelopes in the same m/z region represents the co-eluting peptides with a different charge state. Black triangles indicate the centroid value. Red and blue dotted lines visualize the differences in m/z for the representative peptides of αB peptide and αF helices of HD.

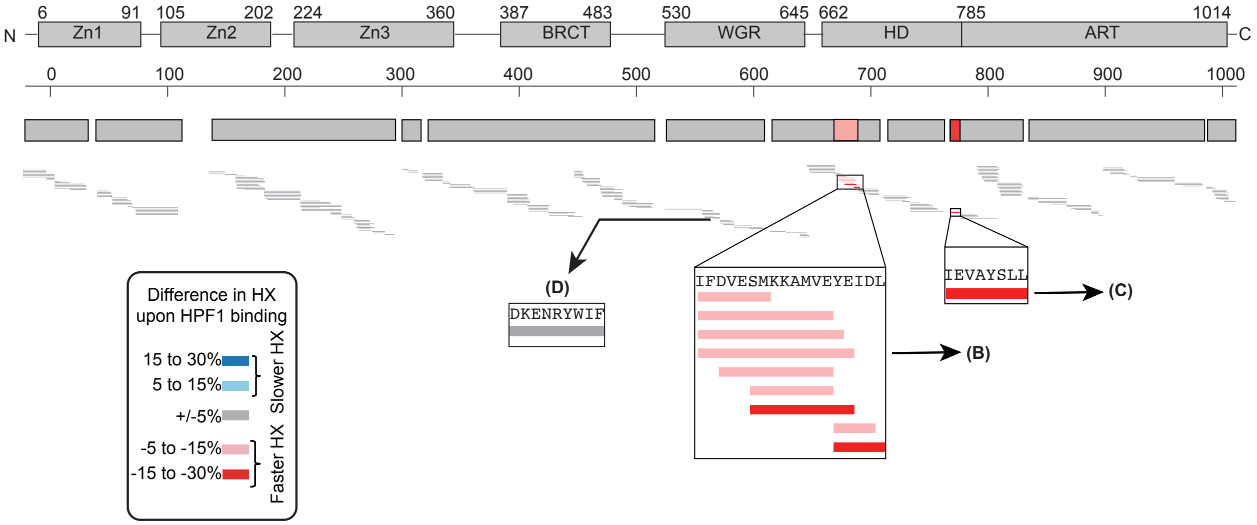


**Supplement Figure 8: HXMS analysis of HPF1 interaction with PARP1 bound to a DNA SSB at 100 s.** The difference plot was obtained by subtracting the percent deuteration of PARP1/DNA/HPF1 complex (at 1:1 HPF1:PARP1) complex from PARP-1/DNA complex at 100 s. Each horizontal bar represents a peptide. The difference plot indicates that most of the peptides (grey) in PARP1/DNA/HPF1 complex have similar HX rates when compared to PARP1/DNA complex. However, peptides in αB and αF helices of HD showed faster exchange (red). The white regions in the difference plot represent gaps in the peptide coverage.
